# Adjusting for systematic technical biases in risk assessment of gene signatures in transcriptomic cancer cohorts

**DOI:** 10.1101/360495

**Authors:** Adrià Caballé Mestres, Antonio Berenguer Llergo, Camille Stephan-Otto Attolini

## Abstract

In recent years, many efforts in clinical and basic research have focused on finding molecular features of tumor samples with prognostic or classification potential. Among these, the association of the expression of gene signatures with survival probability is of special interest given its relatively direct applicability in the clinic and its power to shed insights into the molecular basis of cancer.

Although great efforts have been invested in data processing to control for unknown sources of variability in a gene-wise manner, little is known about the behaviour of gene signatures with respect to the effect of technical variables.

Here we show that the association of signatures with survival may be biased due to technical reasons and propose a simple and low intensive methodology based on correction by expectation under gene randomization. The resulting estimates are centred around zero and ensure correct asymptotic inference. Moreover, our methodology is robust against spurious correlations between global dataset tendencies and clinical outcome.

All tools (will be soon) available in the "HRunbiased" R package as well as processed datasets for colorectal and breast cancer.

---

In recent years, many efforts in clinical and basic research have focused on finding molecular features of tumor samples with prognostic or classification potential [1]. Large cohorts consisting of transcriptional and genetic data have been generated with the aim of characterizing tumor subtypes, understanding biological processes linked to tumorigenesis and identifying molecular profiles associated with clinical parameters [2, 3, 4]. Among these, the prognostic power associated to the expression of single genes or gene signatures has been of special interest, potentially leading to insight into the mechanisms involved in relapse and metastasis, and to the development of prognostic tests for clinical application [5, 6, 7, 8] (Fig 1a).

Summarization of expression data from gene sets to measure a feature of interest is justified by biological and statistical reasons. Studies on large cohorts have shown that pathways are altered through a wide variety of mechanisms, resulting in very low prevalence of single gene alterations [9] (Fig 1b above). As a consequence, transcriptomic analyses at the gene level may suffer from low statistical power and reproducibility [10, 11]. From the statistical perspective, the real status of a pathway is more accurately and sensitively captured by a combination of the expressions of its genes [12] (Fig 1b below). A variety of methods are available for transcriptomic data summarization: average of standardized expression values [12], reduction to a unique component derived from Single Value Decomposition (SVD) and related methods [13], or summaries based on Gene Set Enrichment Analysis (GSEA) [14, 15].

High-throughput data is susceptible to technical variability that can mask true biological information (decrease of statistical power) and/or lead to erroneous conclusions (bias) [16, 17] even if substantial effort has been invested in data processing [18, 19]. For this reason, a number of statistical approaches exist that aim to identify and control for unknown sources of variability while estimating gene expression in a gene-wise manner [18, 20]. Nevertheless, the impact of such effects on gene set summarization may be severe due to possible coordinated effects on all or a large fraction of genes [19, 21]. Therefore, an evaluation of this phenomenon is needed in the specific context of gene set summarization.

In this work, we evaluate two widely used methodologies for pathway summarization: Gene Set Variation Analysis (GSVA) [15] and a z-score based method (ZScore) [12]. Concerns regarding bias and statistical power are explored in a collection of public datasets [22, 23, 24, 25] focusing on cancer prognosis assessment. Drawbacks associated to these methodologies are identified and addressed using a simple strategy. Implications of these findings on conventional statistical inference are discussed, specially concerning the interpretation of asymptotic tests.

**Figure 1:**
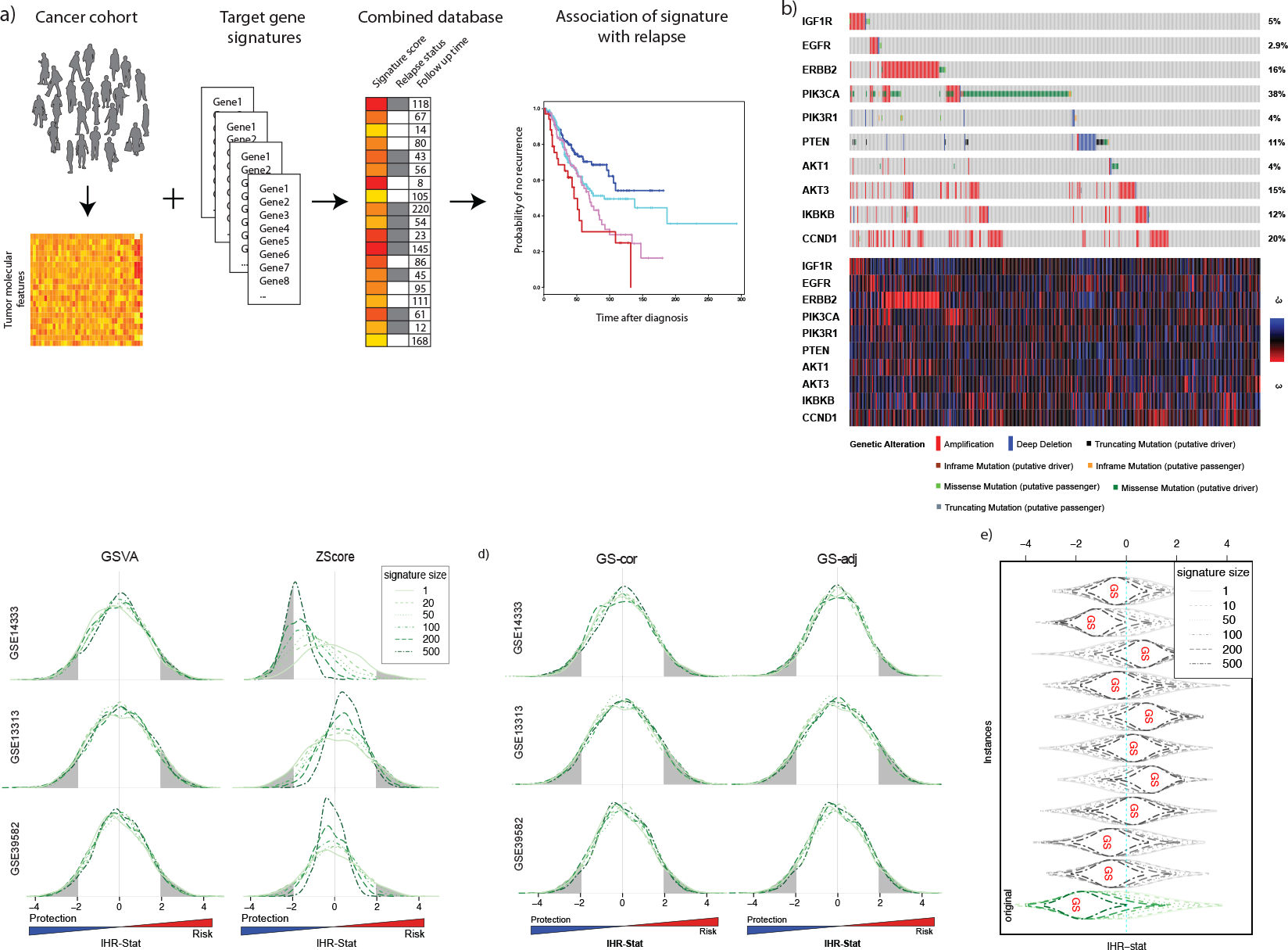
**(a)** Schematic representation of the process to estimate the association of gene signatures and prognosis. **(b)** Genomic alterations (above) in PIK3a pathway genes in TCGA BRCA samples and their corresponding expression (below). **(c)** Density of lHR-stat for random signatures. For ZS-core, all cohorts showed biased means and asymmetric distributions. The proportion of significant tests deviates from the theoretically expected 0.05 and are unequally distributed among protective and risk effects. **(d)** Density of lHR-stat after correction by GS. Both methods of tested result in centred distributions with equal variances for all signature sizes tested. **(e)** Randomization simulations. Relapse status and time were randomized in order to study spurious associations between random signatures and outcome. In all instances, the lHR of random signatures converge towards the lHR of the GS as the signature size increases.

We downloaded and processed 3 independent cohorts of colorectal cancer (CRC) and 6 independent cohorts of breast cancer (see online methods 1), and generated random signatures ranging in size from 1 (single gene) to 500.

For any signature, we applied two strategies for pathway summarization: i) ZScore: gene expression values were converted to z-scores and averaged for each sample, using a similar approach to that in [12]; and ii) GSVA: Gene Set Variation Analysis [15] was applied on each sample and resulting scores were used as summaries. In both cases scores were centred and scaled to make association measures comparable. For each dataset separately, a Cox model was fitted to each score to evaluate their relationship with diseasefree-survival (See online methods 2). The log hazard ratio (lHR) and its associated test statistic (lHR-stat) were used as measures of association. Methods based on SVD represent a variation of the ZScore approach and were not evaluated in this work, as they are prone to be dominated by a small number of genes capturing specific signal which can be different from the biological target (Supp. Fig. 8.1).

For the ZScore method, we found that means of the random lHR deviated from zero in a substantial amount and in different direction depending on the dataset under study, suggesting a strong component of technical origin causing these deviations (Fig 1c, Fig. S1). Moreover, the proportion of random signatures that had significant association with relapse was far from the expected 5% under the hypothesis of lHR = 0 at 5% significance level, compromising the validity and interpretation of statistical inference based on asymptotic assumptions. As the number of genes increased, correlation between random signatures and a Global Signature (GS) including all genes in the dataset quickly converged to 1 (Fig. S3), translating also to convergence in terms of lHRs (Fig. 1e). These results were confirmed in different simulated scenarios and were evident even when only small deviations from zero were observed in average at the gene level (Supp. mat. 3 for permutations-based study). In contrast, GSVA scores were approximately centred in the (a-priori) expected value of zero with similar distributions for all considered sizes (Fig 1c).

These observations motivated the use of the GS to correct the signatures by the expected value if their genes were selected at random, following an analogous strategy to that in [22]. Two approaches were used for GS correction: i) for each sample, we subtracted the Global Signature (GS) values to those obtained by the ZScore method before performing hypothesis testing (GS-cor); and ii) the GS was included as a covariate in the Cox models used for prognosis evaluation (GS-adj). The resulting lHRs of random signatures from GS-cor and GS-adj showed densities centred around zero and with similar variances across all signature sizes (Fig 1d). Moreover, asymptotic inference provided an approximate coverage of 5% under the null hypothesis of zero lHR, and similar to those using GSVA scores (Supp. Tables 3.1-3.6).

Despite the conceptual similarities among GSVA, GS-cor and GS-adj (see online methods 3), one key difference lies in that GS-adj accounts both for the magnitude and the direction of the GS scores and their correlation with the target signature in relation to the outcome by means of the variance-covariance matrix of the model coefficients. To explore this aspect, we compared the summarization methods in different simulated scenarios (See online methods 4)); for type I error, random signatures were categorized according to their correlations with the GS before assessment, while a set of positive control genes (F-TBRS) known to be associated with higher risk in colorectal cancer [23] was used for evaluation of type II error.

For methods based on a-priori correction (GSVA and GS-cor), random signatures produced lHRs distributions shifted from zero in an amount that depended on the correlation between the target signature and the GS (Fig 2a). This dependency was also observed for type II error: while negative correlations increased the chance to detect the association between F-TBRS and relapse, positive correlations considerably decreased statistical power. These results indicate that GSVA and GS-cor are not exempt from biases such as those in Fig. 1b and result in impaired asymptotic inference, possibly due to overcorrection of the global tendency of the dataset. On the contrary lHRs derived from GS-adj showed virtually identical and centred distributions for all correlation levels in the null hypothesis setups (Fig 2a), and only the expected decrease in statistical power was observed as (anti)correlation of GS and F-TBRS increased due to collinearity (Fig 2b). In the real CRC data GS-adj performed equally or better than GS-cor and GSVA in terms of statistical power except for GSE14333 (Fig 2c - Supp. mat. 4); in agreement with the simulation results, this dataset was the only one showing a clear inverse association between the GS and relapse (Fig 1c, Fig 2a) and negative correlation with the F-TBRS (corr = -0.2), which suggests overcorrection of GSVA and GS-cor as an explanation for their apparent higher performance (Supp. mat. 2 for extra simulations).

For settings where the GS is suspected to carry unknown or unidentifiable biological information, we modified the definition of GS by including only low variable genes in its computation (LV-adj), assuming that they are less likely to carry real biological information. For the CRC datasets, we found an increase in statistical power at expense of efficiency in bias correction (Fig 2d). Main surrogate variables or main factors found by the SVA [18] or RUV [20] approaches, respectively, were also considered for the adjusted Cox models without improving the results of LV-adj (Supp. mat. 5).

**Figure 2:**
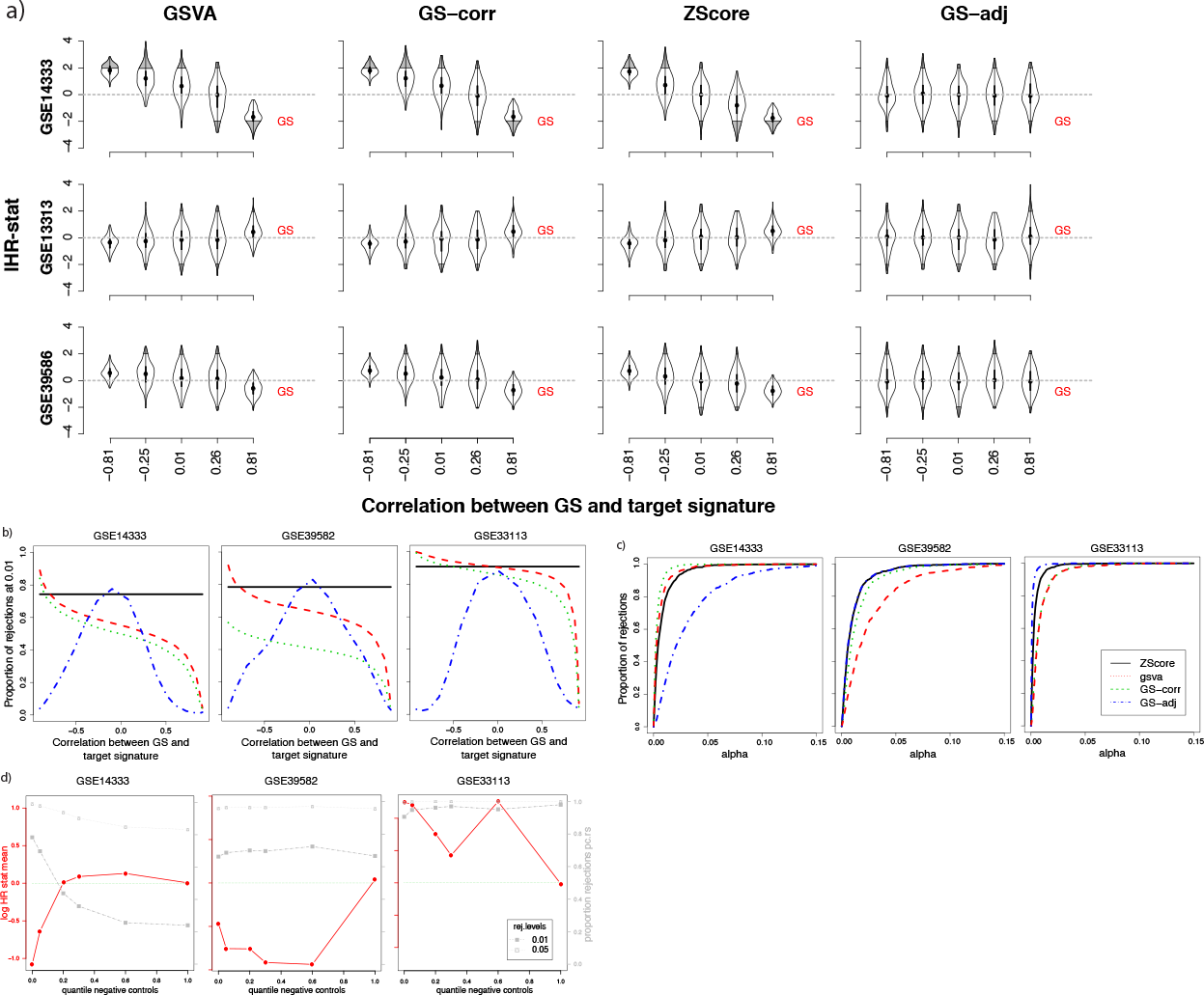
**(a)** The lHR-stat depends on the correlation between GS and target signature. A simulation experiment shows large biases in the lHR for all methods except the GS-adj. These biases depend on the correlation between the GS and the signature of interest. **(b)** Simulation power analysis. In the GSVA and GS-cor methods, negative correlations between F-TBRS and GS increase the chance to detect significant associations between F-TBRS and relapse, while positive correlations harmed considerably their statistical power. GS-adj behaved as expected with maximum power at or near zero correlation between covariables. **(c)** Power analysis in CRC datasets. The power was measured as the proportion of F-TBRS signature rejections at *alpha* significance level. A highly different behaviour was observed in the GSE14333 dataset where the correlation between GS and the F-TBRS signature and the association between GS and the outcome were both negative. **(d)** Correction by low variable genes. Two y-axis plots showing the average lHR-stat for random signatures and the proportion of rejections at *α* = 0.01(0.05) for F-TBRS signatures. The distribution of the lHRs for LVadj random signatures were not completely centred around zero, while the percentages of significant positive controls were slightly higher than those obtained from GS-adj scores, indicating an increase in statistical power at expense of efficiency in bias correction (Tables S5.1-S5.3)

All the analyses in this work were also performed on a selection of BRCA datasets using the MammaPrint [7] signature for prediction of relapse [Supp. mat. 1], which provided analogous conclusions. Although in a lesser degree, issues related to biases and type II error persisted when datasets were combined through merging their expression matrices (Online methods 5 for details) (Supp. mat. 6).

In conclusion, we raise awareness on the usage of gene expression signatures to find associations with survival and prognosis in high-throughput data since, due to the existence of unknown sources of technical variation, the derived estimates may be biased and the corresponding inference inaccurate. Our results show that these biases are driven by overall trends that are present in the gene correlation structure of the expression matrix and by the association of these trends with the outcome, are possibly dataset specific, and quickly increase with the signature size. To address this problem, we propose a simple, flexible and low-intensive computational method (GS-adj) that summarizes the overall data signal to be used as a confounding factor in the statistical analysis. In contrast to existing methodology, our suggested approach accounts not only for the magnitude but also for the correlation of the GS and the signature being evaluated and their association with the outcome. This strategy ensures the validity of standard asymptotic inference for signature testing, and confers advantage in terms of accuracy and interpretability in comparison to other methods based on a-priori correction such as GSVA (Supplementary discussion).

The methods and interpretations derived in this work can be naturally extended to a vast range of domains of high-throughput data such as proteomics, methylation or metabolomics, and many other types of outcomes such as continuous measures (correlation) or binary status (logistic regression). Finally, we provide preprocessed datasets for CRC and BRCA (to be available soon) and a set of tools to diagnose custom datasets and to report adjusted lHR and p-values for different covariates (to be available soon).

## References

[1] Beane, J., Campbell, J. D., Lel, J., Vick, J. & Spira, A. Genomic approaches to accelerate cancer interception. The Lancet Oncology 18, e494–e502 (2017). URL https://www-sciencedirect-com.sire.ub.edu/science/article/pii/S147020451730373X?via

[2] Hutter, C. & Zenklusen, J. C. The Cancer Genome Atlas: Creating Lasting Value beyond Its Data. Cell 173, 283–285 (2018). URL http://www.ncbi.nlm.nih.gov/pubmed/29625045.

[3] Guinney, J. et al. The consensus molecular subtypes of colorectal cancer. Nature Medicine 21, 1350–1356 (2015). URL http://www.nature.com/articles/nm.3967.

[4] Parker, J. S. et al. Supervised risk predictor of breast cancer based on intrinsic subtypes. Journal of clinical oncology: official journal of the American Society of Clinical Oncology 27, 1160–7 (2009). URL http://ascopubs.org/doi/10.1200/JCO.2008.18.1370 http://www.ncbi.nlm.nih.gov/pubmed/19204204 http://www.pubmedcentral.nih.gov/articlerender.fcgi?artid=PMC2667820.

[5] Liu, J. et al. An Integrated TCGA Pan-Cancer Clinical Data Resource to Drive High-Quality Survival Outcome Analytics. Cell 173, 400–416 (2018).

[6] Ferté, C., André, F. & Soria, J.-C. Molecular circuits of solid tumors: prognostic and predictive tools for bedside use. Nature Reviews Clinical Oncology 7, 367–380 (2010). URL http://www.nature.com/articles/nrclinonc.2010.84.

[7] Cardoso, F. et al. 70-Gene Signature as an Aid to Treatment Decisions in Early-Stage Breast Cancer. New England Journal of Medicine 375, 717–729 (2016).

[8] Sanz-Pamplona, R. et al. Clinical Value of Prognosis Gene Expression Signatures in Colorectal Cancer: A Systematic Review. PLoS ONE 7, e48877 (2012).

[9] Bailey, M. H. et al. Comprehensive Characterization of Cancer Driver Genes and Mutations. Cell 173, 371–385.e18 (2018). URL http://www.ncbi.nlm.nih.gov/pubmed/29625053.

[10] Ein-Dor, L., Kela, I., Getz, G., Givol, D. & Domany, E. Outcome signature genes in breast cancer: is there a unique set? Bioinformatics 21, 171–178 (2005).

[11] Ein-Dor, L., Zuk, O. & Domany, E. Thousands of samples are needed to generate a robust gene list for predicting outcome in cancer. Proceedings of the National Academy of Sciences 103, 5923–5928 (2006).

[12] Lee, E., Chuang, H. Y., Kim, J. W., Ideker, T. & Lee, D. Inferring pathway activity toward precise disease classification. PLoS Computational Biology 4 (2008).

[13] Tomfohr, J., Lu, J. & Kepler, T. B. Pathway level analysis of gene expression using singular value decomposition. BMC Bioinformatics 6, 1–11 (2005).

[14] Subramanian, A. et al. Gene set enrichment analysis: A knowledge-based approach for interpreting genome-wide expression profiles. Proceedings of the National Academy of Sciences 102, 15545–15550 (2005). NIHMS150003.

[15] Hanzelmann, S., Castelo, R. & Guinney, J. GSVA: gene set variation analysis for microarray and RNA-Seq data. BMC Bioinformatics 7 (2013).

[16] Scherer, A. Batch effects and noise in microarray experiments: sources and solutions (J. Wiley, 2009).

[17] Bullard, J. H., Purdom, E., Hansen, K. D. & Dudoit, S. Evaluation of statistical methods for normalization and differential expression in mRNA-Seq experiments. BMC Bioinformatics 11, 94 (2010). URL http://bmcbioinformatics.biomedcentral.com/articles/10.1186/1471-2105-11-94.

[18] Leek, J. T. & Storey, J. D. Capturing heterogeneity in gene expression studies by surrogate variable analysis. PLoS Genetics 3, 1724–1735 (2007). NIHMS150003.

[19] Eklund, A. C. & Szallasi, Z. Correction of technical bias in clinical microarray data improves concordance with known biological information. Genome Biology 9, R26 (2008). URL http://genomebiology.biomedcentral.com/articles/10.1186/gb-2008-9-2-r26.

[20] Gagnon-Bartsch, J. A. & Speed, T. P. Using control genes to correct for unwanted variation in microarray data. Biostatistics 13, 539–552 (2012).

[21] Lim, W. K., Wang, K., Lefebvre, C. & Califano, A. Comparative analysis of microarray normalization procedures: effects on reverse engineering gene networks. Bioinformatics 23, i282–i288 (2007).

[22] Efron, B. & Tibshirani, R. On testing the significance of sets of genes. The Annals of Applied Statistics 1, 107–129 (2007). 0610667v2.

[23] Calon, A. et al. Dependency of colorectal cancer on a TGF-beta-driven program in stromal cells for metastasis initiation. Cancer cell 22, 571–584 (2012).

[24] Venet, D., Dumont, J. E. & Detours, V. Most random gene expression signatures are significantly associated with breast cancer outcome. PLoS Computational Biology 7 (2011).

[25] Goeman, J. J. & Buhlmann, P. Analyzing gene expression data in terms of gene sets: methodological issues. Bioinformatics 23, 980–987 (2007). URL https://academic.oup.com/bioinformatics/article-lookup/doi/10.1093/bioinformatics/

